# Simultaneous deep generative modeling and clustering of single cell genomic data

**DOI:** 10.1101/2020.08.17.254730

**Authors:** Qiao Liu, Shengquan Chen, Rui Jiang, Wing Hung Wong

## Abstract

Recent advances in single-cell technologies, including single-cell ATAC-seq (scATAC-seq), have enabled large-scale profiling of the chromatin accessibility landscape at the single cell level. However, the characteristics of scATAC-seq data, including high sparsity and high dimensionality, have greatly complicated the computational analysis. Here, we proposed scDEC, a computational tool for single cell ATAC-seq analysis with deep generative neural networks. scDEC is built on a pair of generative adversarial networks (GANs), and is capable of learning the latent representation and inferring the cell labels, simultaneously. In a series of experiments, scDEC demonstrates superior performance over other tools in scATAC-seq analysis across multiple datasets and experimental settings. In the downstream applications, we demonstrated that the generative power of scDEC helps to infer the trajectory and intermediate state of cells during differentiation and the latent features learned by scDEC can potentially reveal both biological cell types and within-cell-type variations.

---

The organization of chromatin accessibility across the whole genome reflects an epigenetic landscape of gene regulation^1,2^. With the recent development in single-cell technology, it becomes feasible to characterize the epigenetic landscape of individual cells^3^. In particular, single-cell ATAC-seq (scATAC-seq) is an efficient method for the study of variation in chromatin accessibility both between and within populations at single cell level^4,5^. However, the analysis of scATAC-seq presents unique methodological challenges due to the high dimensionality (hundreds of thousands possible peaks) and high data sparsity (only 1-10% peaks are detected per cell) ^6^.

Several computational approaches have been proposed to tackle the challenges in scATAC-seq analysis. scABC estimated weights of cells based on the number of distinct reads and applied a weighted *k*-medoids clustering to infer cell types^7^. cisTopic applied latent Dirichlet allocation (LDA) as a probabilistic model to identify the *cis*-regulatory topics enriched in different cells by optimizing topic-cell probability and region-topic probability simultaneously^8^. Cusanovich et al. proposed a pipeline which performs the term frequency-inverse document frequency transformation (TF-IDF) and singular value decomposition (SVD) iteratively to get a low dimensional representation of scATAC-seq data^4,9^. Scasat introduced another pipeline which involved Jaccard similarity measure and multidimensional scaling (MDS) to reduce the high dimensionality in scATAC data^10^. SnapATAC divided genome into bins with equal size and builds a bins-by-cells binary count matrix and then applied principle component analysis (PCA) for a dimension reduction^11^. Recently, deep generative models have emerged as a powerful framework for both representation learning and data generation^12–14^. A newly developed method SCALE utilized a variational autoencoder (VAE) to learn the latent features of scATAC-seq data and then used a *K*-means by default for clustering the latent features^15^.

Here, we proposed a new approach for analyzing **<u>sc</u>**ATAC-seq data by simultaneously learning the **<u>D</u>**eep **<u>E</u>**mbedding and **<u>C</u>**lustering of the cells in an unsupervised manner. Our method, named scDEC, was based on learning a pair of generative adversarial networks (GANs) (Fig. 1). Such a symmetrical and paired GAN architecture has been recently successfully applied to image style transfer^16^ and density estimation^17^. Here, we adopted this architecture to unsupervised clustering and applied it to the analysis of single cell genomic data. Unlike all current methods discussed above, where an external method (e.g., *K*-means) is typically required for clustering the latent features, in our method the cell clustering process is directly modeled by neural networks. Thus, cell clustering and latent feature representation learning will be jointly optimized during the training process. In other words, scDEC enables simultaneous learning of low dimensional embedding and cell clustering. We demonstrate the advantage of this approach in a series of experiments, where scDEC is shown to outperform other current methods. We also illustrated several downstream applications of scDEC in scATAC-seq analysis, including trajectory inference, donor effect removal and latent feature interpretation.

**Fig. 1.**
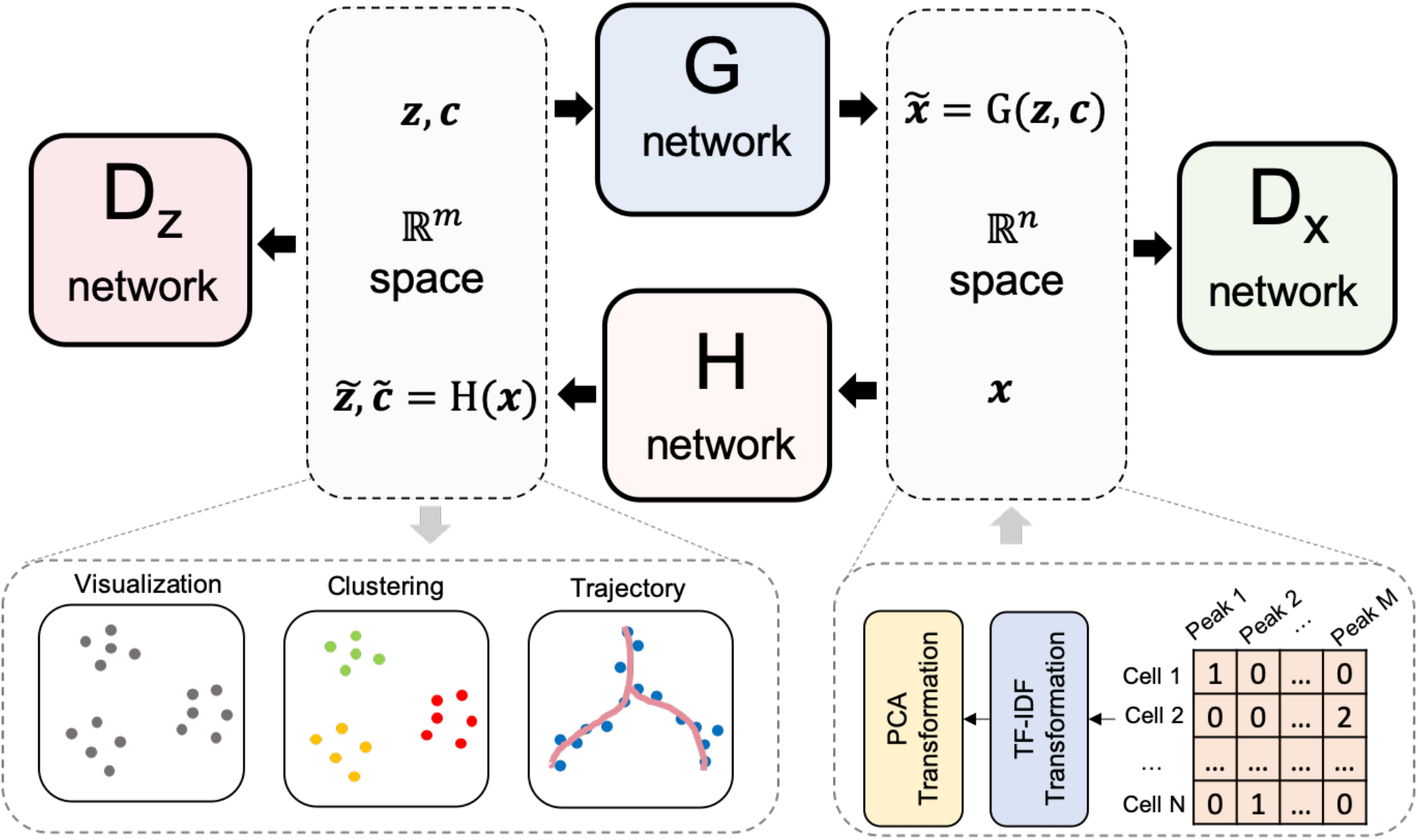
The illustration of scDEC model. The read count matrix of scATAC-seq will first be preprocessed by a TF-IDF transformation and a PCA dimension reduction (e.g., *n*=20) before it is fed to the scDEC model. In the latent space, latent variables ***z*** and ***c*** sampled from a Gaussian distribution and a Category distribution respectively, will be concatenated together before they are fed to the G network. The H network has two outputs of which one corresponds to the latent embedding 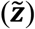 and one corresponds to the estimated cluster label 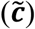 through a softmax function. The D_*x*_ network works as a discriminator for discerning the true scATAC-seq data (***x***) from the generated data 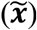. The D_*z*_ network is another discriminator for distinguishing the learned continuous latent variable 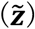 from the real continuous latent variable (***z***).

## Results

### Overview of scDEC model

scDEC consists of two GAN models, which are utilized for transformations between latent space and data space (Fig. 1). The scATAC-seq data will first be preprocessed through TF-IDF transformation and a PCA dimension reduction before it is fed to the scDEC model. Assume the input scATAC-seq data contains *K* cell types, a continuous latent variable ***z*** and a discrete latent variable ***c*** will be introduced, where 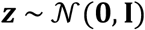 and **c** ~ Cat(*K*, ***w***), respectively. We also provided an approach for estimating the number of cell subpopulations if *K* is unknown (Methods). The forward transformation through the G network can be considered as a process of conditional generation given an encoded style (***z***) and an indicated cluster label (***c***). The backward transformation through the H network aims at encoding a data point ***x*** to the latent space and inferring the cluster label, simultaneously. If we assume the last layer of H network contains *m* nodes (*m*>*K*), then 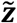 denotes the output of the first (*m*-*K*) nodes and 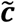 denotes the output of the remaining *K* nodes with an additional softmax function. D_*x*_ and D_*z*_ are two discriminator networks which are used for matching the distributions of data 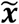 and 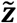 to the empirical distribution of the data and latent variable distribution, respectively. (G, D_*x*_) and (H, D_*z*_) can be considered as two GAN models that are jointly trained. The G and H network each contains 10 fully-connected layers while D_*x*_ and D_*z*_ each has two fully-connected layers (see detailed hyperparameters in Supplementary Table 1). Note that the weights ***w*** in the Category distribution is also learned automatically via an updating scheme according to the feedback of inferred cluster labels by 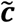 (Methods). After model training, the cluster label can be inferred based on 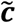 (Methods). The output of the last layer of H network combined with 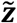 and 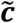 (before softmax) are useful for downstream analysis such as data visualization and trajectory analysis.

### scDEC automatically identifies cell types in scATAC-seq data

To demonstrate the ability of scDEC for revealing differences between different cell subpopulations and identifying cell types in an unsupervised manner, we test it on four benchmark scATAC-seq datasets across different number of cells and cell types (Supplementary Figure 1 and Table 2). Specifically, scDEC was benchmarked against six current methods, including scABC^7^, SCALE^15^, cisTopic^8^, Cusanovich2018^4,9^, Scasat^10^ and SnapATAC^11^ (Methods). The performance of a method is evaluated on 1) whether different cell subpopulations can be clearly separated in a low-dimensional space, and 2) whether true cell type labels can be accurately inferred by clustering. To address the first question, we first applied each method to conduct a dimension reduction or to extract the latent features. The latent dimension is set to 15 for the two datasets with relatively smaller number of cells and cell populations, and 20 for the two larger datasets. For each method, we constructed a t-SNE^18^ or UMAP^19^ plot based on the latent features and then visualized the truth cell labels on the plot to see whether the subpopulations were well separated. To address the second question, for each method we compare its clustering results to the true subpopulations based on three commonly used metrics, namely Normalized Mutual Information (NMI), Adjusted Rand Index (ARI) and Homogeneity score (Homogeneity) (Methods). The results are summarized below. We note that scDEC’s performance is not sensitive to the dimension of latent space (Supplementary Figure 2)

#### InSilico dataset^5^

This dataset is an in silico mixture constructed by artificially combining six individual scATAC-seq experiments which were separately conducted on a different cell line. We observed that cells from a minor cell type TF-1 (6.83%, in purple) were dispersed into several clusters by SCALE, Cusanovich2018, Scasat and SnapATAC while cisTopic and scDEC can well maintain the close distance in the low-dimensional representation (Fig. 2a). scDEC achieves an NMI of 0.871, an ARI of 0.896, and a Homogeneity of 0.866, which outperforms the best baseline method scABC (NMI=0.822, AIR=0.855, and Homogeneity=0.840) by a large margin (Fig. 2e and Supplementary Figure 3).

**Fig. 2.**
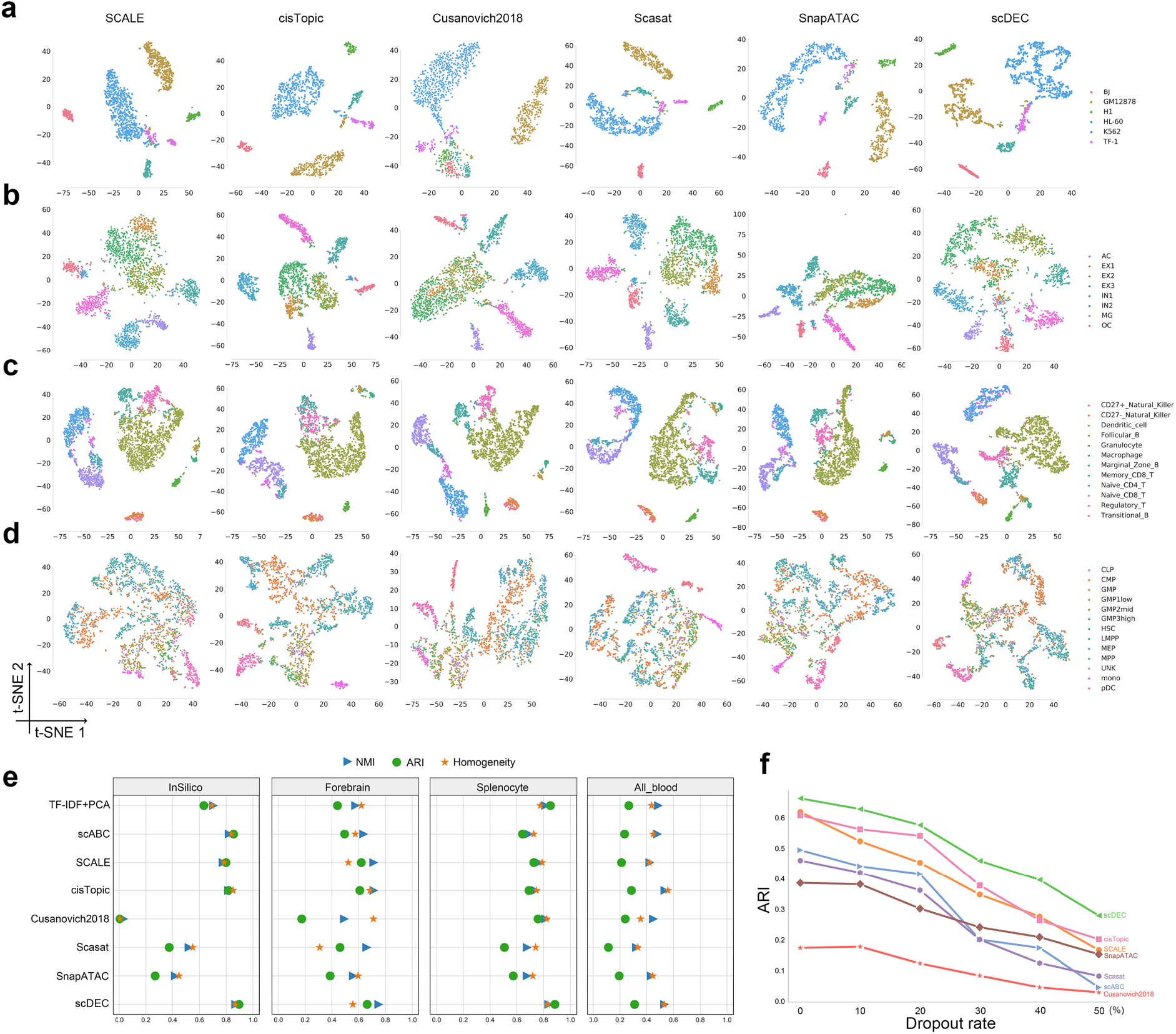
Evaluation of scDEC compared with other baseline methods. **a**. Visualization of InSilico dataset by different methods. **b**. Visualization of Forebrain dataset by different methods. **c**. Visualization of Splenocyte dataset by different methods. **d**. Visualization of All_blood dataset by different methods. **e**. Clustering results of different methods across four datasets. **f**. Performance of different methods under different dropout rate on the Forebrain dataset.

#### Forebrain dataset^20^

This dataset was derived from P56 mouse forebrain cells which contained eight different cell groups in adult mouse forebrain. Interestingly, all the baseline methods failed to distinguish three subtypes of excitatory neuron cells (EX1, EX2 and EX3) while scDEC showed a relatively clear separation among these three subpopulations of cells (Fig. 2b). Again, scDEC demonstrates a superior clustering performance by achieving the highest NMI of 0.750 and ARI of 0.663 (Fig. 2e and Supplementary Figure 4).

#### Splenocyte dataset^21^

This dataset was collected from a mixture of mouse splenocytes after removing red blood cells, which finally resulted in 12 cell subpopulations. A major cell type follicular B cells (42.89%, in brown) and two subtypes of CD8 cells are more or less mixed together with other subpopulations of cells by all baseline methods while scDEC illustrates a clearer separation (Fig. 2c). As the largest dataset (around 3k cells) among the four datasets, scDEC still achieves the highest NMI of 0.839, ARI of 0.884 and Homogeneity of 0.829 (Fig. 2e and Supplementary Figure 5).

#### All blood dataset^22^

This dataset involves cellular differentiation of multipotent cells during human hematopoiesis, containing 13 subpopulations of cells in total. Three types of cells, including monocyte cells (mono), plasmacytoid dendritic cells (pDC) and CLP cells, can only be separated from other cells by cisTopic, Scasat and scDEC (Fig. 2d). scDEC ranks first in both ARI (0.309) and Homogeneity (0.531) and ranks second in ARI (0.533), which is slightly lower than cisTopic (ARI=0.536) (Fig. 2e and Supplementary Figure 6).

Next, we further investigated the performance of different methods at different dropout rate, in order to assess the ability of handing scATAC-seq data with different degree of sparsity. We downsampled the original reads in the Forebrain dataset by randomly dropped out the non-zero entities in the read count matrix with probability equal to the dropout rate. scDEC consistently demonstrates the best performance *w.r.t* the ARI metric for clustering at different dropout rate ranging from 0 to 50%. At the dropout rate of 50%, scDEC achieves an ARI of 0.279, compared to 0.202 of the best baseline cisTopic (Fig. 2f).

### scDEC facilitates cell type-specific motif discovery and trajectory inference

We next explored whether scDEC can help identity cell-type specific motifs, which is essential for understanding the context-specific gene regulation. To achieve this, we first applied scDEC model to the mouse forebrain dataset^20^ to infer the cluster label for each individual cell, and used chromVAR^23^ to identify cluster-specific enriched motifs from the JASPAR database^24^. We then ranked cluster-specific enriched motifs (Methods) and discovered several significant motif enrichment patterns (Fig. 3a, Supplementary Table 3). We observed both single cluster-specific motifs and the co-occurrence of motifs in two (cluster 1 and 6) or three clusters (cluster 2,3 and 4), which might reveal the co-regulation mechanism underlying the corresponding multiple TFs. For example, En1, which is enriched in cluster 1, is a well-known marker for the brain fate in astrocytes (AC)^25^. It was reported that Neurod2 regulates the cortical projection neuron which constitute the major excitatory neuron (EX) population^26^. Meis1 was proved to reveal crucial functions in neural differentiation from neural progenitors^27^. Vax1 is a novel homeobox-containing gene that regulates the development of the basal forebrain^28^. The impact of Elk1 deficiency was proved to indicate the microglial (MG) activation^29^. The compound loss of Sox9 will lead to a further decrease in oligodendrocyte (OC) progenitors^30^. These literature-validated motifs were demonstrated in the t-SNE visualization according to the enrichment score calculated by chromVAR (Fig. 3b).

**Fig. 3.**
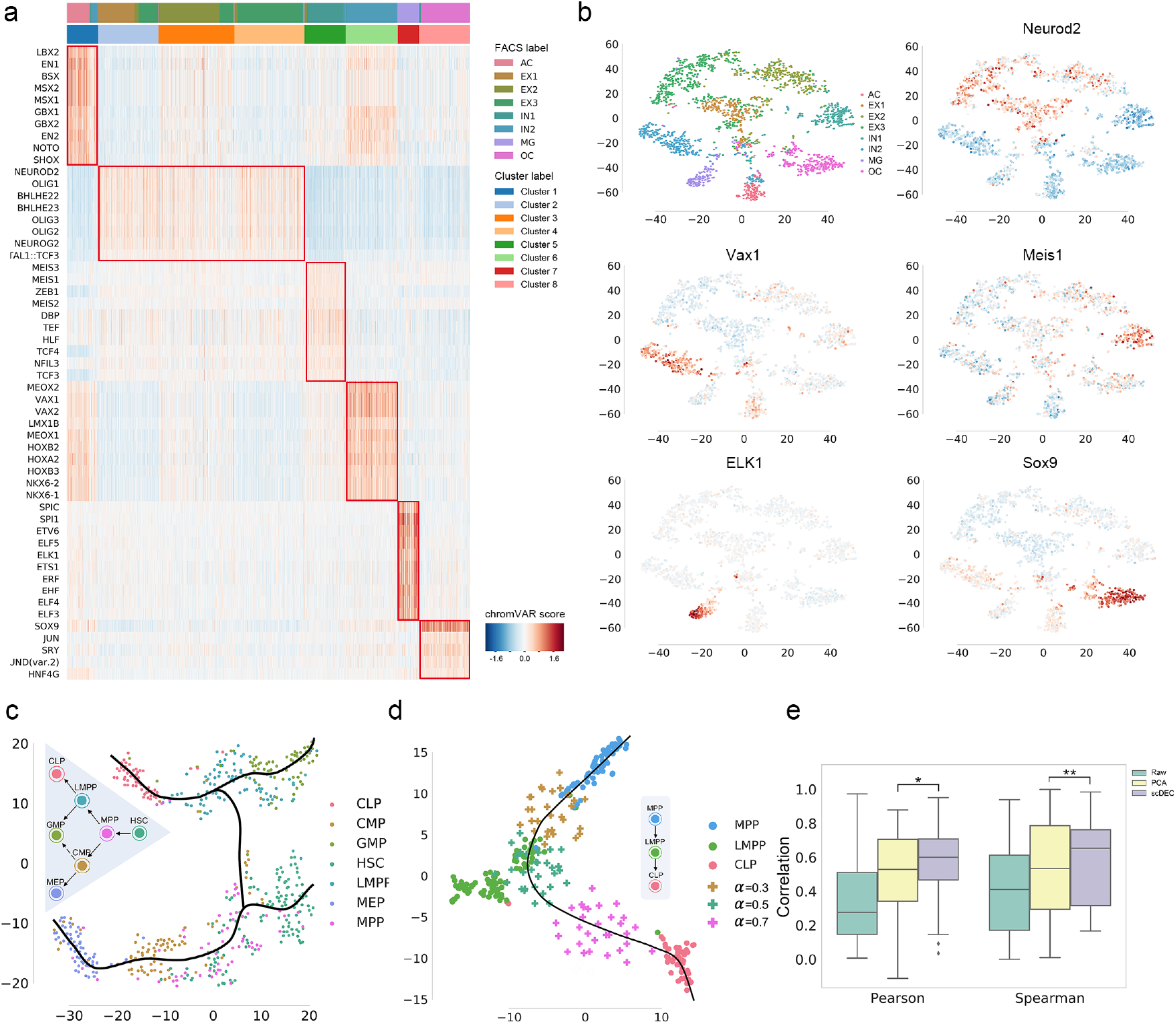
Cluster-specific motif recovery and trajectory inference. **a**. Heatmap of enriched motifs, each row denotes a motif and each column denotes a cell. Both cluster label and FACS label were provided and aligned. **b**. The t-SNE visualization of several literature-validated motifs. **c**. The hematopoiesis differentiation trajectory inferred by scDEC. **d**. The generated intermediate state between MPP and CLP. 30 data points were generated at different generation coefficient α. **e**. The generated intermediate scATAC data by interpolation on the latent label indicator has a higher correlation with the meta cell (the average profile of ground truth cells) than the scATAC-seq that were directly interpolated on the raw data and PCA reduced data. * *p*-value<1.28×10^−16^, ** *p*-value< 4.40×10^−8^

Next, we applied scDEC to trajectory inference during the hematopoiesis differentiation. We collected the cells from the donor BM0828 of the All blood dataset, which contains 533 cells across 7 subpopulations at different stage of differentiation. After obtaining the low-dimensional representation and t-SNE projection of scATAC-seq data, we can annotate the smooth curves which represent different cell lineages with the help of Slingshot software^31^ (Fig. 3c). The smooth curves with a tree-based structure are largely consistent with the true hematopoietic differentiation tree. Although it has been proved that CMP can differentiate into both GMP and MEP^32^, only differentiation path from CMP to MEP was observed in this dataset. We then took the cells from MPP, LMPP and CLP for a further study, where there exists a differentiation path (MPP→LMPP→CLP). To fully exploit the generation power of scDEC, we first left LMPP out as the target cells for imputation and trained scDEC based on the remaining cells composing of only MPP and CLP cells. Then we imputed data by interpolating the latent label indicator (Methods) and visualized the imputed data together with the true data. Interestingly, when the interpolation coefficient α changes from 0 to 1, the imputed data seem to capture the dynamics differentiation path from MPP to CLP. Specifically, the generated scATAC-seq data are similar to the real LMPP data according to t-SNE visualization when α = 0.5 (Fig. 3d). Next, we asked whether the interpolation on the latent indicator is a more effective way of data generation than directly interpolating on the raw scATAC-seq. We averaged all the scATAC-seq data of LMPP cells as a meta-cell and calculated the Pearson correlation between generated data and meta-cell. The generated data by scDEC achieves a significantly higher correlation than generated data by direct interpolation and interpolation on PCA reduced data (Fig. 3e and Supplementary Table4). To sum up, the generation power of scDEC shed light on recovering the missing cell types of scATAC data and exploring the intermediate state of two neighboring cell types of scATAC-seq data.

### scDEC disentangles donor effect and promotes interpretation of latent features

Single-cell experiments are often conducted with notable differences in capturing time, equipment and even technology platforms, which may introduce batch effects in the data. To evaluate whether scDEC can automatically correct or alleviate batch effect in the training process. We collected human hematopoietic cells containing three cell types (CLP, LMPP and MPP) from two donors BM0828 and BM1077^22^. We mixed the cells from two donors together and evaluated how well the variation due to cell types and donors are resolved in the embedding (i.e., latent representation) learned by scDEC and alternative methods. Note that the latent dimension of each method was fixed to 13 and no donor information was revealed to each method. Since the embedding by scDEC depends on the number of clusters *K*, we varied *K* from 2 to 6 and examine the gap statistic plot (Fig. 4d), which exhibited two peaks at *K*=3 and *K*=5, respectively. The embedding results for scDEC and alternative methods were shown in Fig. 4a and Supplementary Figure 7-8. It is seen that the three cell types as well as the donor effects in two of the cell types are well captured by scDEC (*K*=5), cisTopics and SnapATAC, but not by SCALE, whereas the donor effect in the third cell type (CLP) is too small to be discernible. It is interesting that at *K*=3 (the first peak of the gap statistic) the clustering results by scDEC matches the three cell types almost perfectly. Specifically, SCALE is basically unable to separate the three type of cells clearly. cisTopic and SnapATAC cannot alleviate the donor effect in LMPP or MPP cells as the same type of cells from two different donors were separated with a notable distance in the t-SNE plot (Fig. 4a). Considering the first mode where *K*=3, only 9 cells from donor BM0828 and 17 cells from donor BM1077 were wrongly clustered by scDEC, which illustrates a total error rate of 6.86%. Besides, scDEC also demonstrates an NMI of 0.754, ARI of 0.805 and Homogeneity of 0.757 which outperforms other comparing methods by a large margin (Fig. 4b and Supplementary Figure 8). In this sense our method can be used to adjust for donor- or batch-effects in clustering and visualization.

**Fig. 4.**
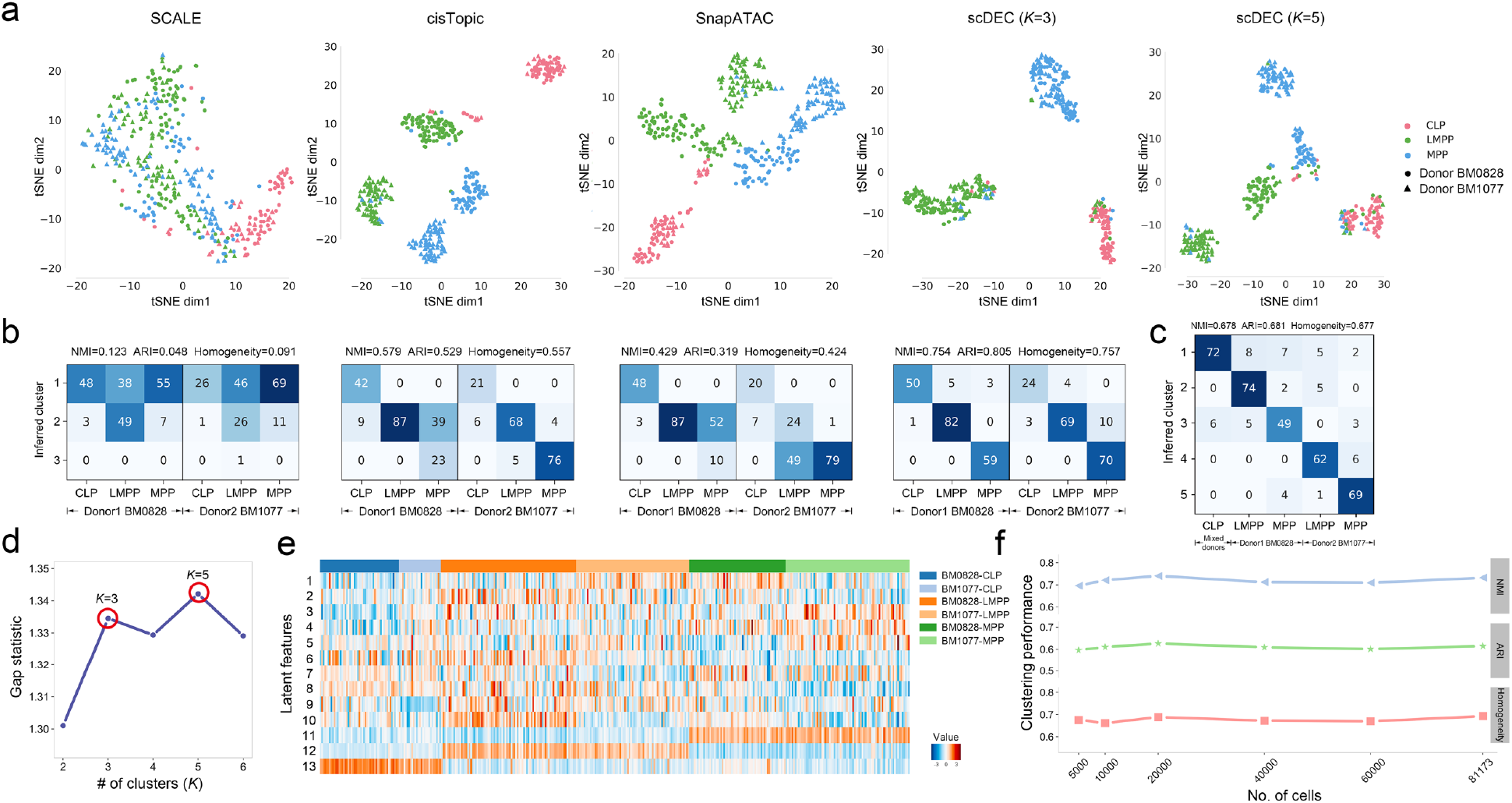
scDEC alleviates donor effect and is robust to large dataset. **a**. The t-SNE visualization of the latent features learned by different methods. Different colors denote different cell types and different shape (circle or triangle) represents which donor it comes from. For scDEC, different *K* (3 and 5) results in different latent features visualization. **b**. The confusion matrix of the clustering by scDEC and comparing methods (*K*=3). The NMI, ARI and Homogeneity are also annotated on the top of the confusion matrix. **c**. The confusion matrix of the clustering by scDEC when *K*=5. **d**. The gap statistic shows two modes at *K*=3 and *K*=5, respectively. **e**. The visualization of the latent features learned by scDEC. The first 10 dimensions correspond to the continuous latent variable 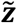 and the last three features correspond to the discrete latent variable 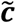. **f**. The clustering performance of scDEC when applying to a large mouse atlas dataset.

Next, we carefully analyzed the latent feature learned by scDEC by visualization. We noticed that features corresponding to the latent discrete variable (feature 11-13) were highly correlated to biological cell types while other features more or less revealed within-cell-type variations (Fig. 4e). For example, feature 1 is highly expressed in the donor BM1077 of LMPP and BM0828 of MPP. Feature 10 can be a donor-specific indicator of LMPP. To sum up, the interpretable features in the latent space reveal both biological cell types and within-cell-type variations.

### scDEC is capable of analyzing large scATAC-seq data

We further examined that whether scDEC is applicable to extremely large scATAC-seq dataset. We collected a dataset from a mouse atlas study which contains 81,173 single cells from 13 adult mouse tissues using sci-ATAC-seq^9^. The original atlas study applies a computational pipeline to infer 40 cell types, which were regarded as “reference” cell label for the comparison of scDEC and other baselines methods. To investigate the scalability of scDEC, we randomly down-sampled the original dataset to different scale of dataset and scDEC shows a consistently good agreement with the reference cell label (Fig. 4f). For the full scale of the dataset, scDEC achieves an NMI of 0.732, ARI of 0.614 and Homogeneity of 0.693 while most previous methods failed to handle the full dataset according to a benchmark study^6^. We compared scDEC to the deep learning method SCALE and noticed that scDEC achieves a higher consistency with “reference” label but a little slower running time (Supplementary Figure 9). We also noticed that the scDEC successfully identified most of the major reference cell type for each tissue (Supplementary Figure 10).

## Discussion

In this study, we proposed scDEC for accurately characterizing cell subpopulations in scATAC-seq data using a deep generative model. Unlike previous studies that take dimension reduction and clustering as two independent tasks. scDEC intrinsically integrates the low-dimensional representation learning and unsupervised clustering together by carefully designing a GAN-based symmetrical architecture. scDEC can serve as a powerful tool for scATAC-seq data analysis, including visualization, clustering and trajectory analysis. In a series experiments, scDEC achieves competitive or superior performance compared to other baseline methods. In the downstream applications, we focused on the generation power of scDEC, which can facilitate the intermediate cell state inference. The latent features learned by scDEC reveals both biological cell types and within-cell-type variations, which shed light on helping better understand the biological mechanism.

We also provide several directions for improving scDEC. First, scDEC serves as a general-purpose unsupervised learning framework, which can also be explored for applying to scRNA-seq data or the joint analysis of scRNA-seq and scATAC-seq data. Second, the way of latent indicator interpolation in the data generation can be further explored, especially in a complicated tree or graph-based trajectory of cell differentiation. One more essential problem will be how to use scDEC to generate missing data at one time point given a time-course scATAC-seq datasets.

To sum up, with scDEC, researchers could perform a scATAC-seq analysis of the cell types or tissues with interests. Then, one can simultaneously infer the cell label and uncover the biological findings underlying the learned latent features. We hope scDEC could help unveil the single-cell regulatory mechanism and contribute to understanding heterogeneous cell populations.

## Methods

### Data preprocessing

All the scATAC-seq datasets were uniformly preprocessed before fed to scDEC model. To reduce the level of noise, we only kept peaks that have at least one read count in more than 3% of the cells. Next, similar to Cusanovich et al^9^, we applied a term frequency-inverse document frequency (TF-IDF) transformation to the raw scATAC-seq count matrix, which is widely used technology in information retrieval and text mining^33,34^. It helps increase proportionally to the number of times a peak appears in the cell, which gives a higher importance weight to the peaks with less frequency. Finally, a principle component analysis^35^ (PCA) will be applied to reduce the dimension of the scATAC to 20, which is implemented with “scikit-learn” package^36^. scDEC shows robustness to the dimension of PCA (Supplementary Figure 2). The summary of all scATAC-seq datasets used in this study were provided in Supplementary Table 2.

### Visualization

We use t-distributed stochastic neighbor embedding^18^ (t-SNE) as the default algorithm for visualization the latent features of scATAC-seq data learned by different methods by setting the visualization dimension to 2. The t-SNE was implemented with “Scikit-learn” package^36^. The uniform manifold approximation and projection (UMAP)^19^ was also implemented as an additional visualization tool for latent features.

### Adversarial training in scDEC model

The scDEC model consists a pair of two GAN models. For the forward GAN mapping, G network aims at conditionally generating samples 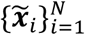 that have a similar distribution to the observation data 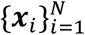 while the discriminator D_*x*_ tries to discern observation data (positive) from generated samples (negative). The backward mapping function H and the discriminator D_*z*_ aims to transform the data from data space to the latent space. Discriminators can be considered as binary classifiers where an input data point will be asserted to be positive (1) or negative (0). We use WGAN-GP^37^ as the architecture for the GAN implementation where the gradient penalty of discriminators will be considered as an additional loss terms. We define the objective loss functions of the above four neural networks (G, H, D_*x*_ and D_*z*_) in the training process as the following

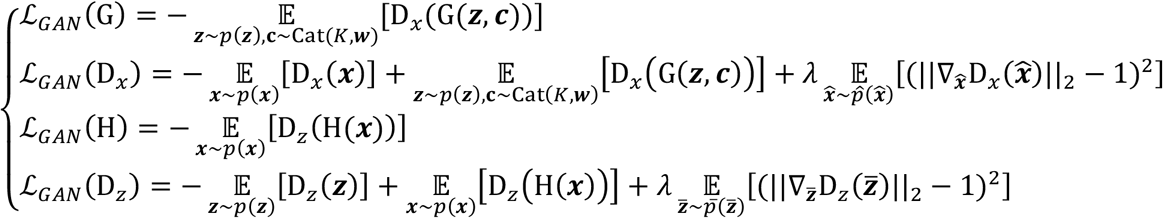

where *p*(***z***) and Cat(*K*, ***w***) denote the probability distribution of continuous variable and discrete variable in the latent space, respectively. In practice, sampling ***x*** from *p*(***x***) can be regarded as a procedure of randomly sampling from *i.i.d* observations data with replacement. 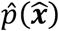 and 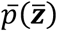 denote uniformly sampling from the straight line between the points sampled from true data and generated data. Minimizing the loss of a generator (e.g., 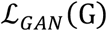) and the corresponding discriminator (e.g., 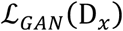) are somehow contradictory as the two networks (G and D_*x*_) compete with each other during the training process. *λ* is a penalty coefficient which is set to 10 in all experiments.

### Roundtrip loss

During the training, we also aim to minimize the roundtrip loss which is defined as ρ((***z***, ***c***), H(G(***z***, ***c***))))and ρ(***x***, G(H(***x***))) where ***z*** and ***c*** are sampled from the distribution of the continuous latent variable *p*(***z***) and the Category distribution Cat(*K*, ***w***). The principle is to minimize the distance when a data point goes through a roundtrip transformation between two data domains. In practice, we used *l*_2_ loss as the continuous part in roundtrip loss and used cross entropy loss as the discrete part in roundtrip loss. We further denoted the roundtrip loss as

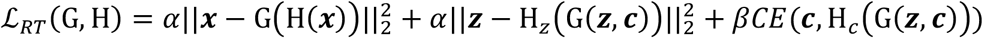

where *α* and *β* are two constant coefficients which are both set to 10. H_*z*_(·) and H_*c*_(·) denote the continuous and discrete part of output from H(·), respectively and *CE*(·) represents the cross-entropy loss function. The idea of roundtrip loss which exploits transitivity for regularizing structured data has also been used in previous works^16,38^.

### Full training loss

Combining the adversarial training loss and roundtrip loss together, we can get the full training loss for generator networks and discriminator networks as 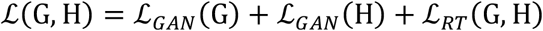 and 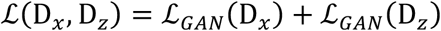, respectively. To achieve joint training of the two GAN models, we iteratively updated the parameters in the two generative models (G and H) and the two discriminative models (D_*x*_ and D_*z*_), respectively. Thus, the overall iterative optimization problem can be represented as

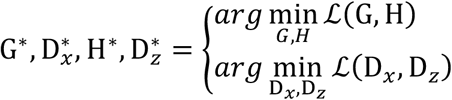

An Adam optimizer^39^ with a learning rate of 2 × 10^−4^ was used for updating the weights in the neural networks. The training process is illustrated in Supplementary Table 5 in details.

### Data generation in scDEC

We generate the state of intermediate cell by interpolating the latent indicator ***c*** of two “neighboring” cell types. Assume there are two cell types which correspond to the latent indicator ***c***_**1**_ and ***c***_**2**_, respectively. The generated data can be represented as 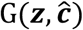 where 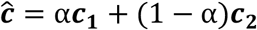. Note that the α is the generation coefficient from 0 to 1 and ***z*** is still sampled from a standard Gaussian distribution. The interpolation of latent features have already been used for exploring and visualizing the transition from two type of images^40^.

### Network architecture in scDEC

All the networks in scDEC are made of fully-connected layers. The G network contains 10 fully-connected layers and each hidden layer has 512 nodes while the H network contains 10 fully-connected layers and each hidden layer has 256 nodes. D_*x*_ and D_*z*_ both contain 2 fully-connected layers and 256 nodes in the hidden layer. Batch normalization^41^ was used in discriminator networks. All hyperparameters were also provided in Supplementary Table 1.

### Updating the Category distribution

The probability ***w*** in the Category distribution Cat(*K*, ***w***) is adaptively updated every 100 batches of data based on the inferred cluster label from 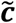 of full training data (Table 1).

**Table 1.**
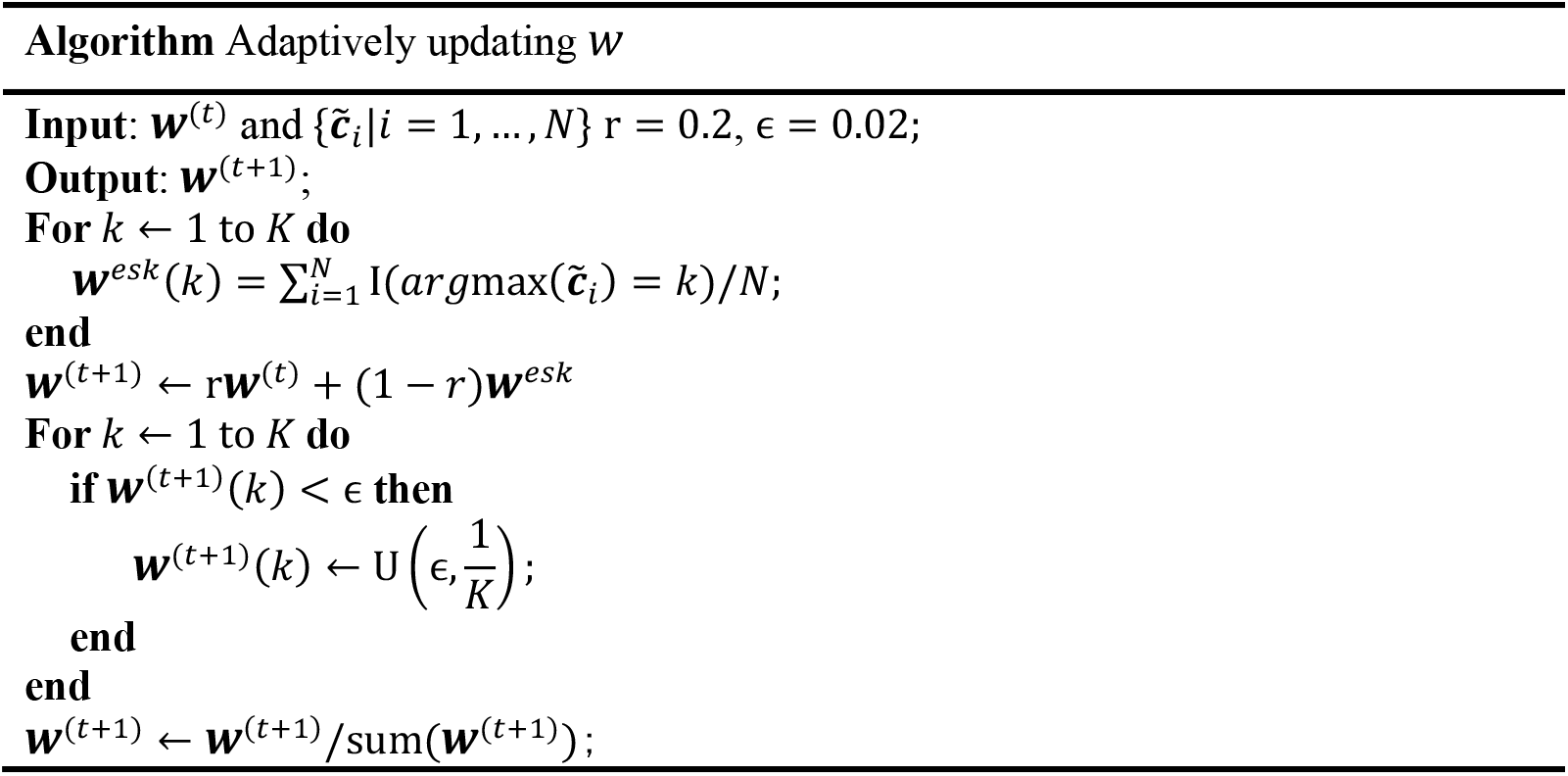
Adaptively updating *w* in the Category distribution in the latent space of scDEC. *r* is the ratio coefficient and ∊ denotes the lower bound of the cluster proportion. U(p, q) represents a uniform distribution between p and q.

### Evaluation metrics for clustering

We compared different methods for clustering according to three metrics, normalized mutual information (NM)I^42^, adjusted Rand index (ARI)^43^ and Homogeneity^44^. Assuming *U* and *V* are true label assignment and predicted label assignment given *n* data points, which have *C*_*U*_ and *C*_*V*_ clusters in total, respectively. NMI is then calculated by

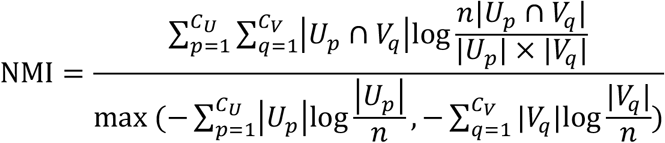

The Rand index^45^ is a measure of agreement between two cluster assignments while ARI corrects lacking a constant value when the cluster assignments are selected randomly. We define the following four quantities 1) *n*_1_: number of pairs of two objects in the same groups in both *U* and *V*, 2) *n*_2_: number of pairs of two objects in different groups in both *U* and *V*, 3) *n*_3_: number of pairs of two objects in the same group of *U* but different group in *V*, 4) *n*_4_: number of pairs of two objects in the same group of *V* but different group in U. Then ARI is calculated by

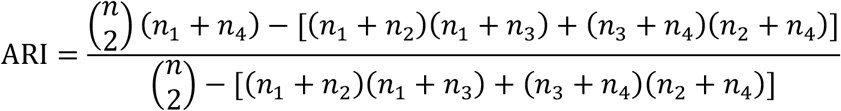

Homogeneity is calculated by 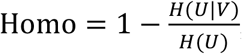, where

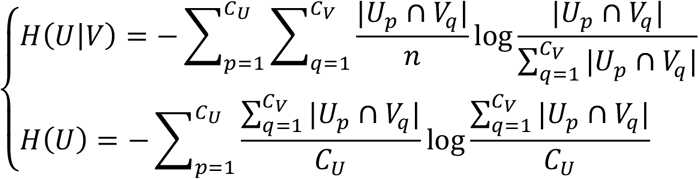

### Estimating number of clusters *K*

In order to apply scDEC to scATAC-seq where the number of cell types is unknown. We provide an algorithm for estimating the number of clusters *K* using gap statistic^46^. We first compared the average within-cluster distance of the preprocessed scATAC-seq data and a reference dataset, which can be constructed with random matrix with the same size using *K*-means algorithm. The average within-cluster distance on the reference dataset was calculated for 1000 times by Monto Carlo simulation and the average result was used. The optimal choice of *K* is given for which the gap between the single cell data and the reference data is maximum. We note that this estimation of number of clusters *K* well matches the truth clusters numbers with the scATAC-seq used in this study (Supplementary Figure 11).

### Identification of cluster-specific motifs

The cluster-specific motifs are identified by Mann-Whitney U test^47^ with the alternative hypothesis that the chromVAR scores^23^ of cells in one cluster or multiple clusters have a positive shift compared with chromVAR scores of the rest of cells. Then the motifs will be ranked according to the *p*-values and the top-ranked motifs were illustrated.

### Baseline methods

We compared scDEC to multiple baseline methods in this study, including scABC^7^, SCALE^15^, cisTopic^8^, Scasat^10^, Cusanovich2018^4,9^ and SnapATAC^11^. The source code for the implementation of baseline methods was downloaded from a benchmark study^6^.

## Data availability

InSilico dataset was collected from GEO database with accession number GSE65360. The mouse forebrain dataset was downloaded from GEO database with accession number GSE100033. Splenocyte dataset can be accessed at ArrayExpress database with accession number E-MTAB-6714. All blood dataset can be accessed at GEO database with accession number GSE96772. The mouse atlas data is available at http://atlas.gs.washington.edu/mouse-atac. All the processed data for the input of scDEC can also be downloaded from https://zenodo.org/record/3984189#.XzDpJRNKhTY.

## Code availability

scDEC is an open-source software based on the TensorFlow library^48^, which can be freely downloaded from https://github.com/kimmo1019/scDEC.

## Acknowledgement

We thank Fengling Chen for her helpful discussion. This work was supported by NIH grants R01 HG010359 and P50 HG007735. This work was also supported by the National Key Research and Development Program of China (No. 2018YFC0910404), the National Natural Science Foundation of China (Nos. 61873141, 61721003, 61573207).

