## Supplementary Materials for "Simultaneous deep generative modeling and clustering of single cell genomic data"

[1]MOE key Laboratory of Bioinformatics, Bioinformatics Division, Beijing National Research Center for Information Science and Technology, Department of Automation, Tsinghua University, Beijing 100084, China; [2]Department of Statistics, Stanford University, Stanford, CA 94305, USA; [3]Department of Biomedical Data Science, Bio-X Program, Center for Personal Dynamic Regulomes, Stanford University, Stanford, CA 94305, USA.

* Corresponding authors:

### Contents

**Supplementary Figures** .......... 3

**Supplementary Tables** .......... 14

### Supplementary Figures

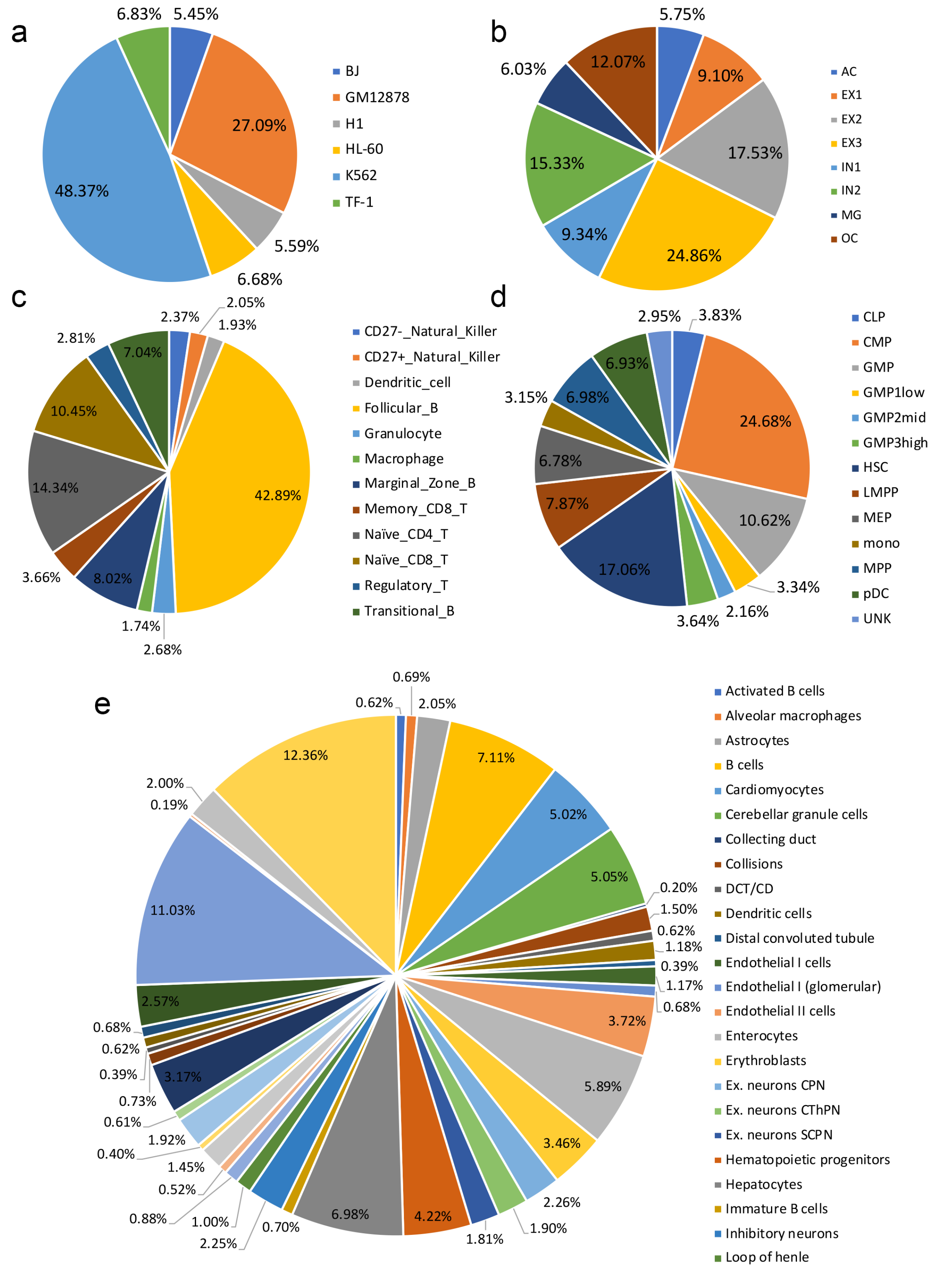


**Supplementary Figure 1.** The statistics of five scATAC-seq datasets used in this study. **a**. InSilico datasets. **b**. Forebrain datasets. **c**. Splenocyte datasets. **c**. All blood datasets. **e**. Mouse atlas dataset.

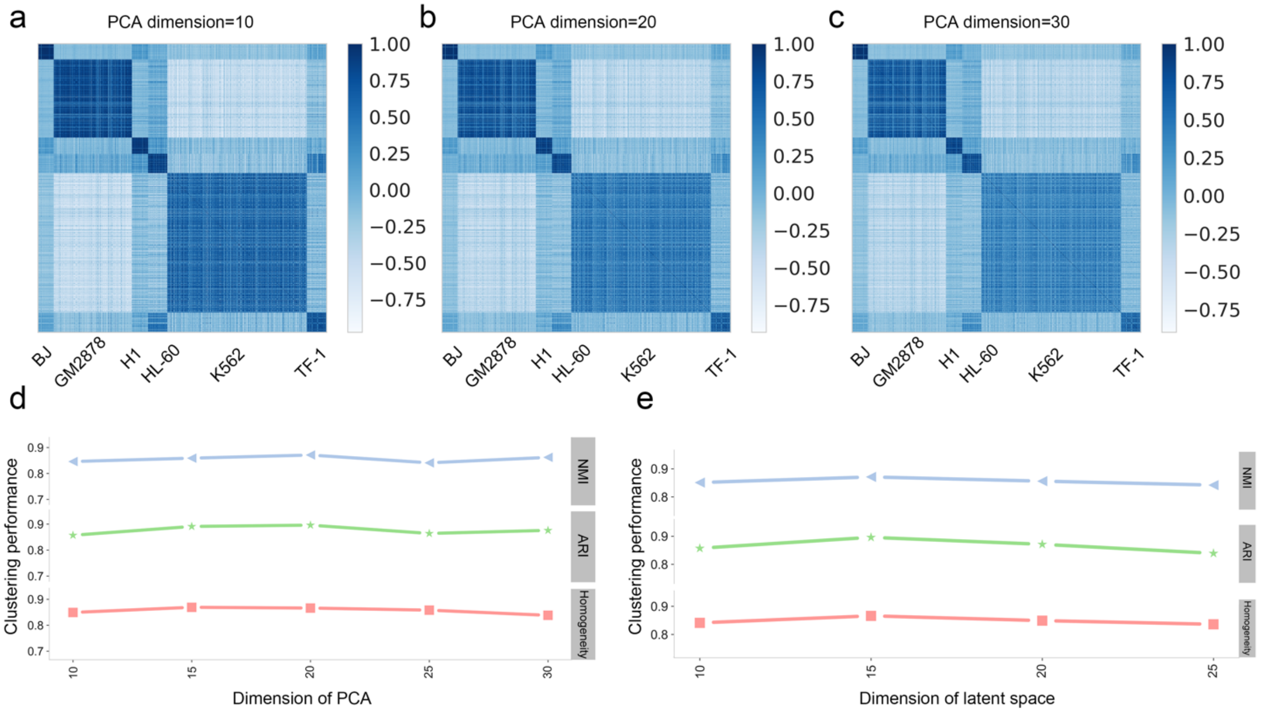


**Supplementary Figure 2.** scDEC model demonstrates a robustness to the dimension of both latent space and the PCA dimension. Taking the Silico dataset for an example. **a**-**c**. We first use different dimension for PCA transformation when preprocessing data. Then cell-to-cell Cosine similarity matrix was calculated. The boundary remains clear when dimension changes from 10 to 30. **d**. The performance of clustering when changing the dimension of PCA from 10 to 30. Note that the latent dimension was fixed to 15.**e**. The performance of clustering when changing the dimension of latent features from 10 to 25. Note that the dimension of PCA was fixed to 20.

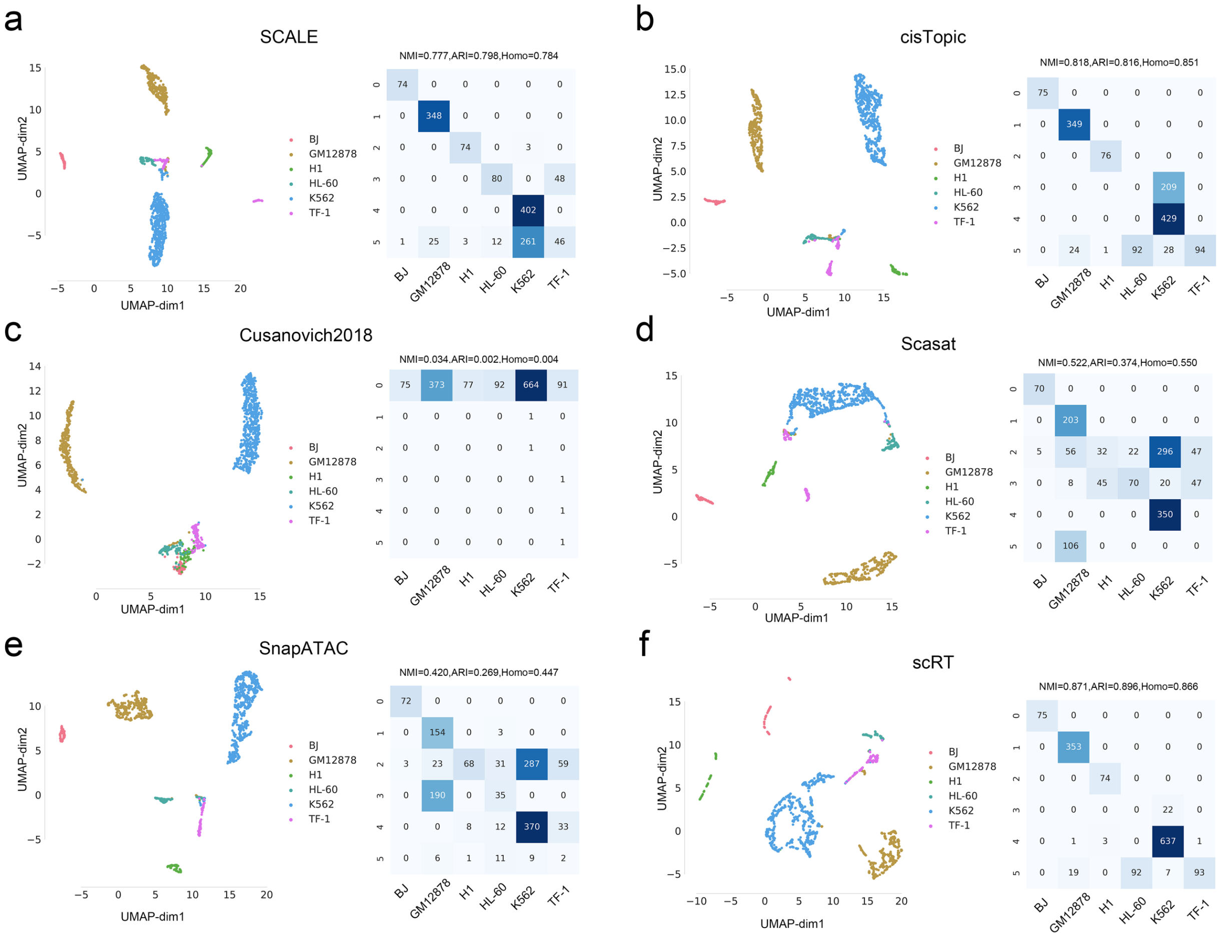


**Supplementary Figure 3.** UMAP visualization and clustering confusion matrix of different methods with InSilico dataset. The NMI, ARI and Homogeneity were shown on the top of each confusion matrix. **a**. SCALE. **b**. cisTopic. **c**. Cusanovich2018. **d**. Scasat. **e**. SnapATAC. **f**. scDEC.

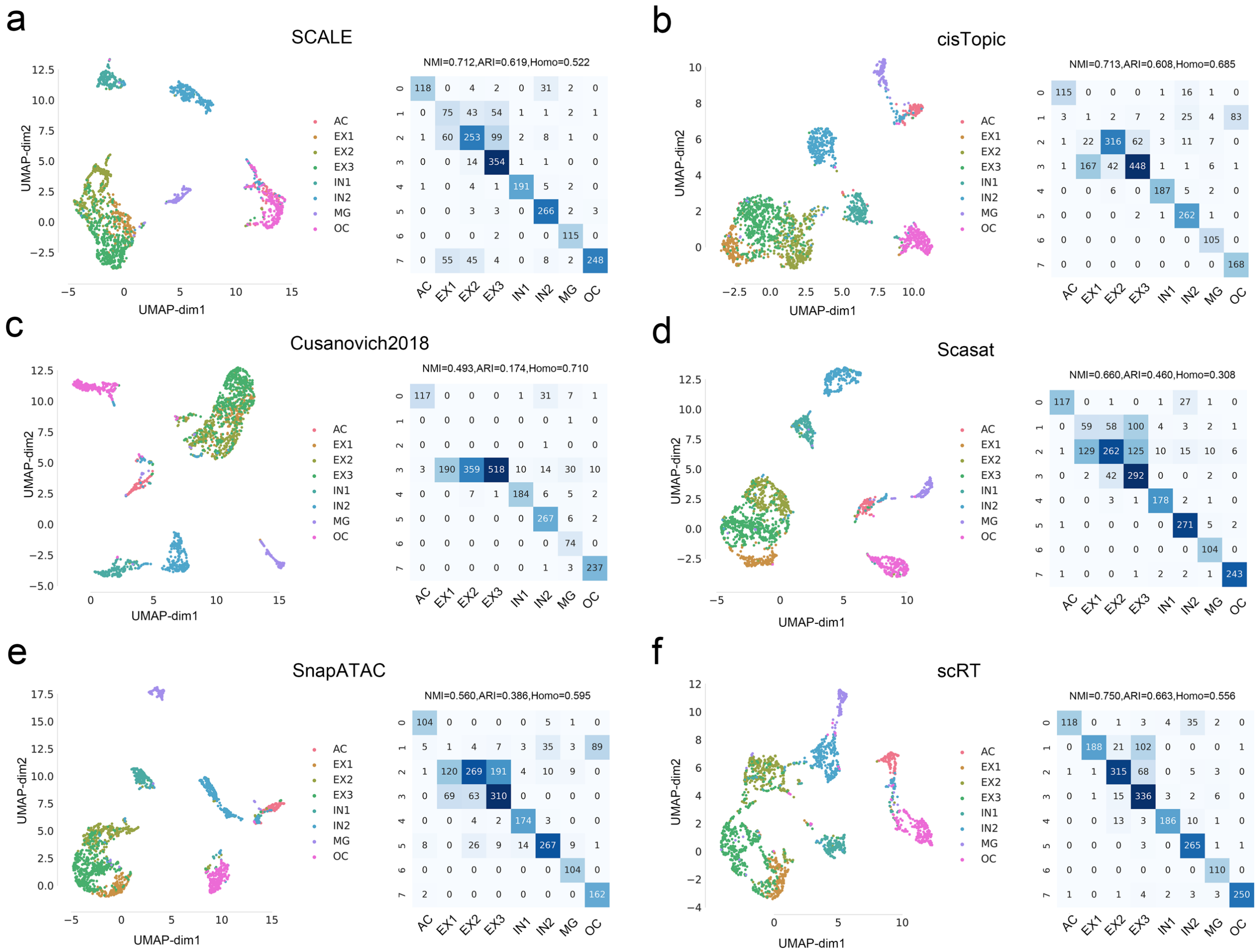

**Supplementary Figure 4.** UMAP visualization and clustering confusion matrix of different methods with Forebrain dataset. The NMI, ARI and Homogeneity were shown on the top of each confusion matrix. **a**. SCALE. **b**. cisTopic. **c**. Cusanovich2018. **d**. Scasat. **e**. SnapATAC. **f**. scDEC.

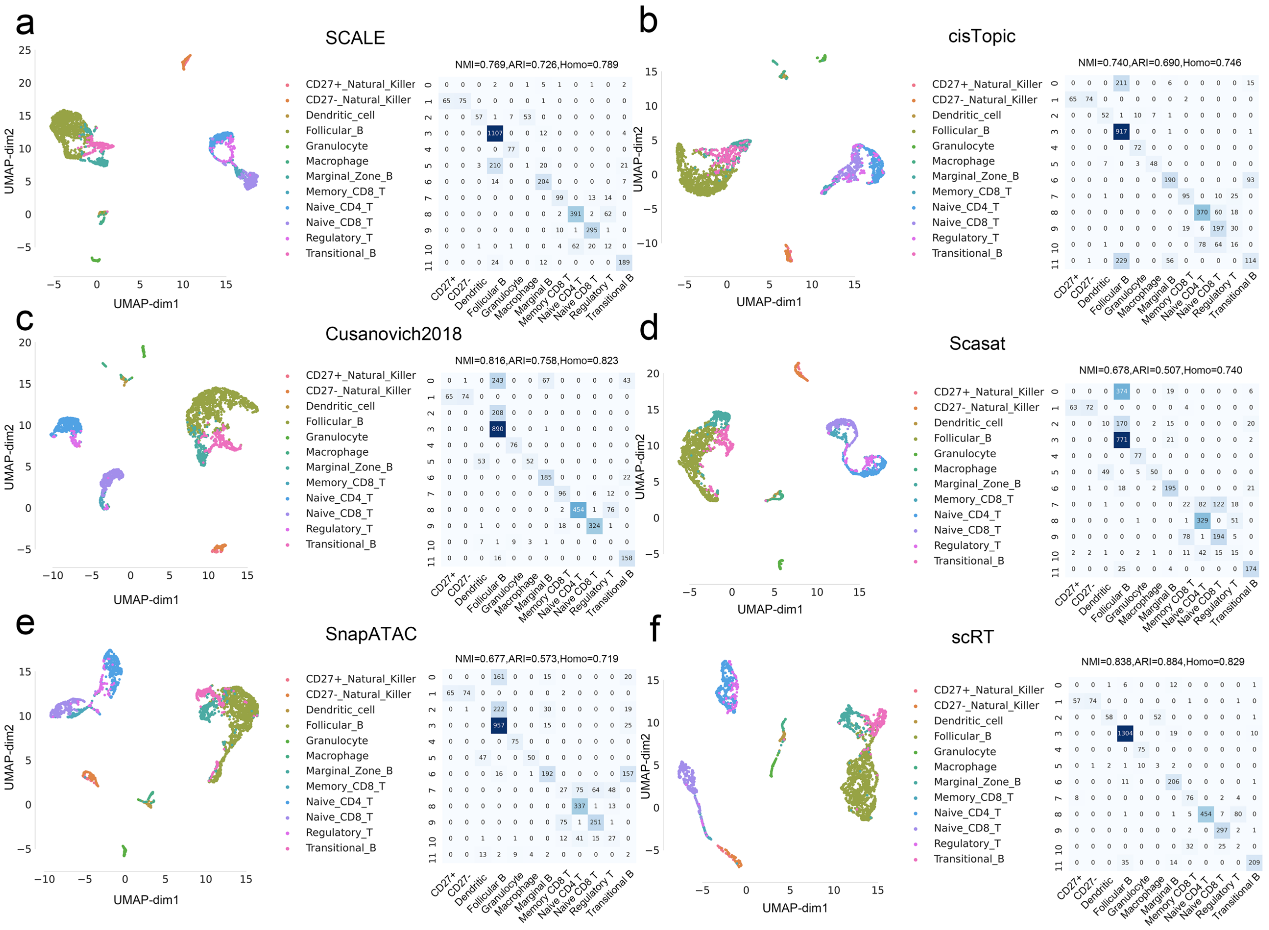


**Supplementary Figure 5.** UMAP visualization and clustering confusion matrix of different methods with Splenocyte dataset. The NMI, ARI and Homogeneity were shown on the top of each confusion matrix. **a**. SCALE. **b**. cisTopic. **c**. Cusanovich2018. **d**. Scasat. **e**. SnapATAC. **f**. scDEC.

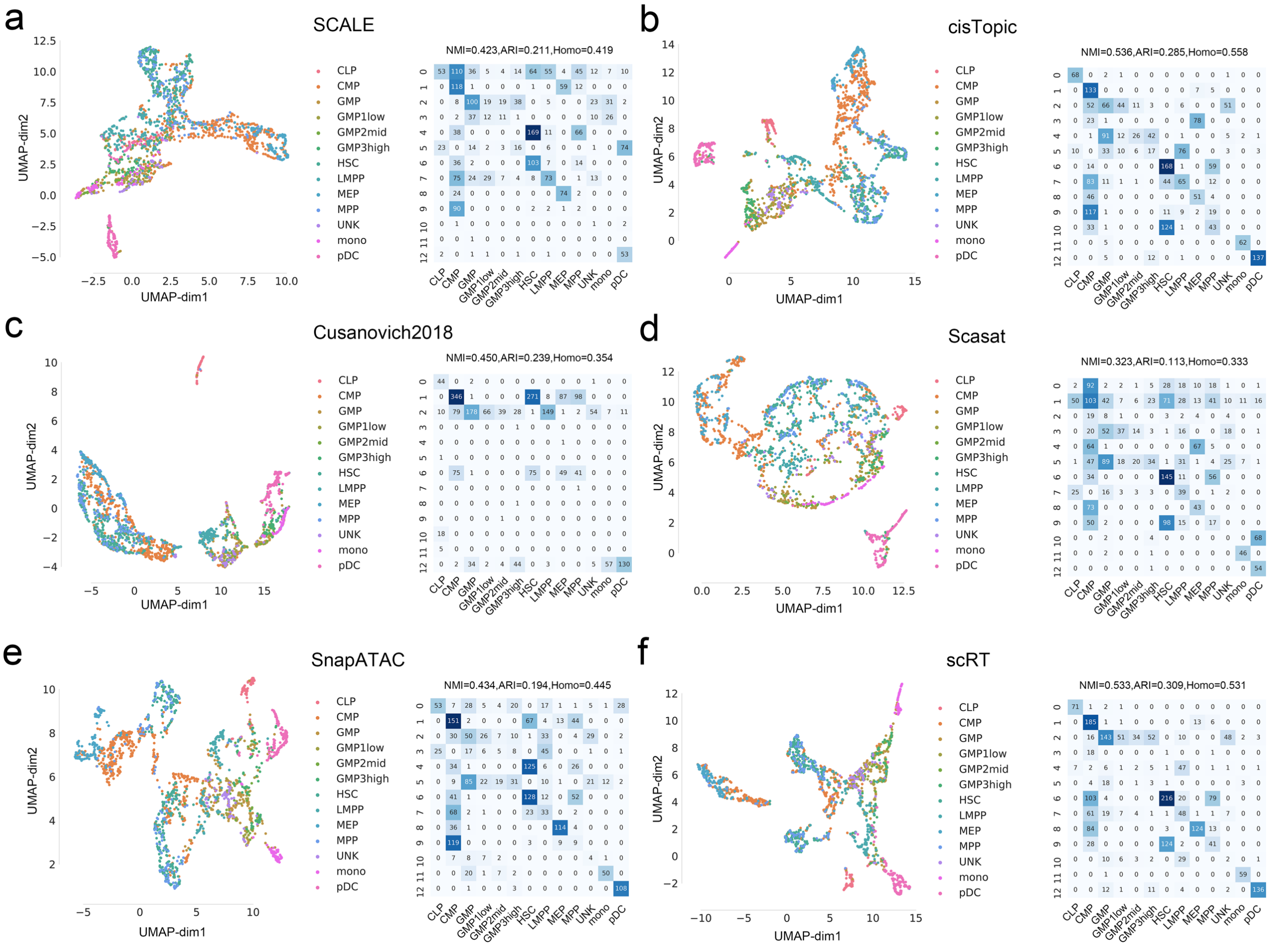

**Supplementary Figure 6.** UMAP visualization and clustering confusion matrix of different methods with All blood dataset. The NMI, ARI and Homogeneity were shown on the top of each confusion matrix. **a**. SCALE. **b**. cisTopic. **c**. Cusanovich2018. **d**. Scasat. **e**. SnapATAC. **f**. scDEC.

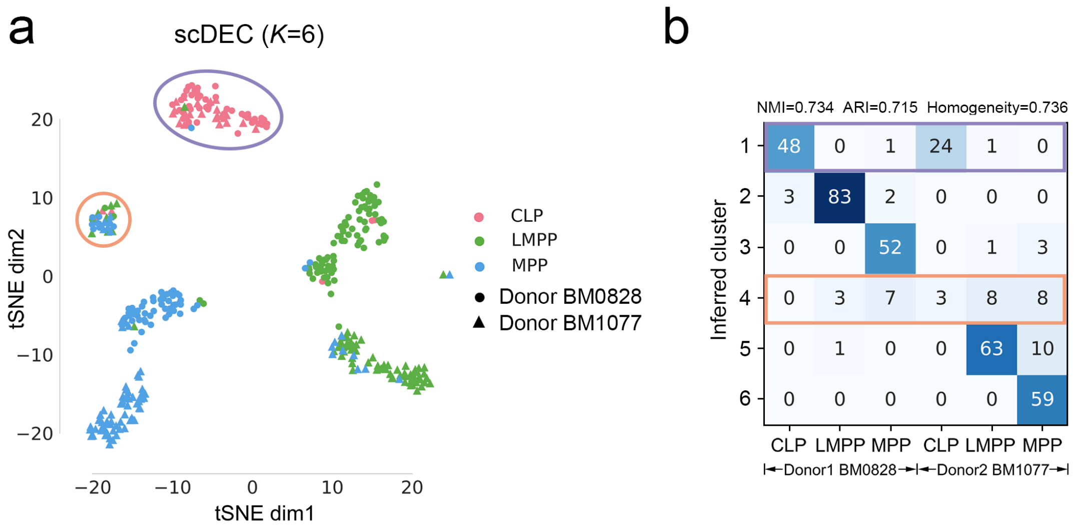


| Inferred cluster | CLP | LMPP | MPP | CLP | LMPP | MPP |
|---|---|---|---|---|---|---|
| 1 | 48 | 0 | 1 | 24 | 1 | 0 |
| 2 | 3 | 83 | 2 | 0 | 0 | 0 |
| 3 | 0 | 0 | 52 | 0 | 1 | 3 |
| 4 | 0 | 3 | 7 | 3 | 8 | 8 |
| 5 | 0 | 1 | 0 | 0 | 63 | 10 |
| 6 | 0 | 0 | 0 | 0 | 0 | 59 |

**Supplementary Figure 7. a.** The t-SNE visualization of scDEC of which *K* is set to 6. As the CLP cells have a small donor variation which cannot be discernable, the CLP cells from two donors were considered as one cluster. So scDEC was forced to automatically learns a newly small cluster (cluster 4). **b**. The confusion matrix of scDEC clustering results. It is clear to see cluster 2, 3, 5 and 6 corresponds to a specific cell type from a specific donor, cluster 1 corresponds to CLP cells from both two donors and cluster 4 is the newly learned small cluster.

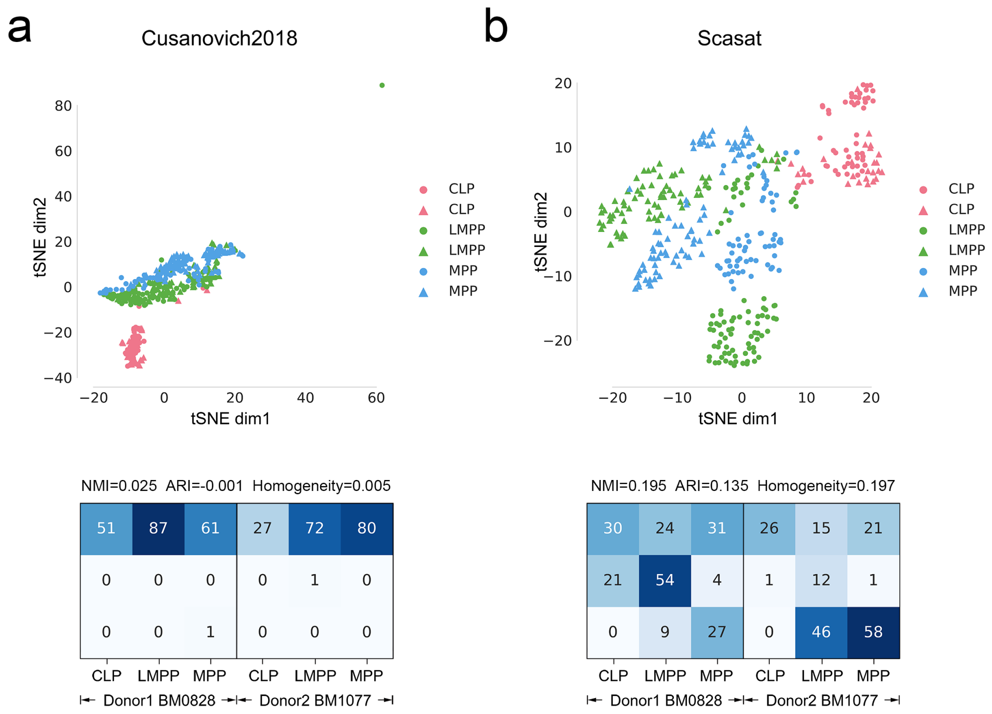


**Supplementary Figure 8.** T-SNE visualization and clustering confusion matrix of Cusanovich2018 and Scasat with CLP, LMPP and MPP cells from two donors. The NMI, ARI and Homogeneity were shown on the top of each confusion matrix. ($K$=3) **a**. Cusanovich2018. **b**. Scasat.

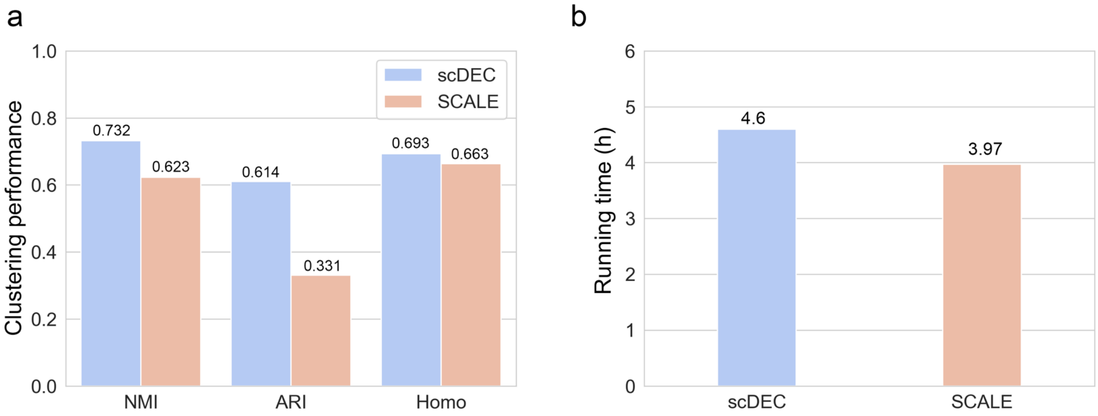


**Supplementary Figure 9.** Mouse atlas full dataset compared to SCALE. **a**. scDEC achieves a more consistent results with the "reference cell label". **b**. scDEC consumes more running time than SCALE with the full mouse atlas dataset. All the experiments were carried on a Linux platform with 2 Intel Xeon E5-2600 CPUs and 512 GB memory.

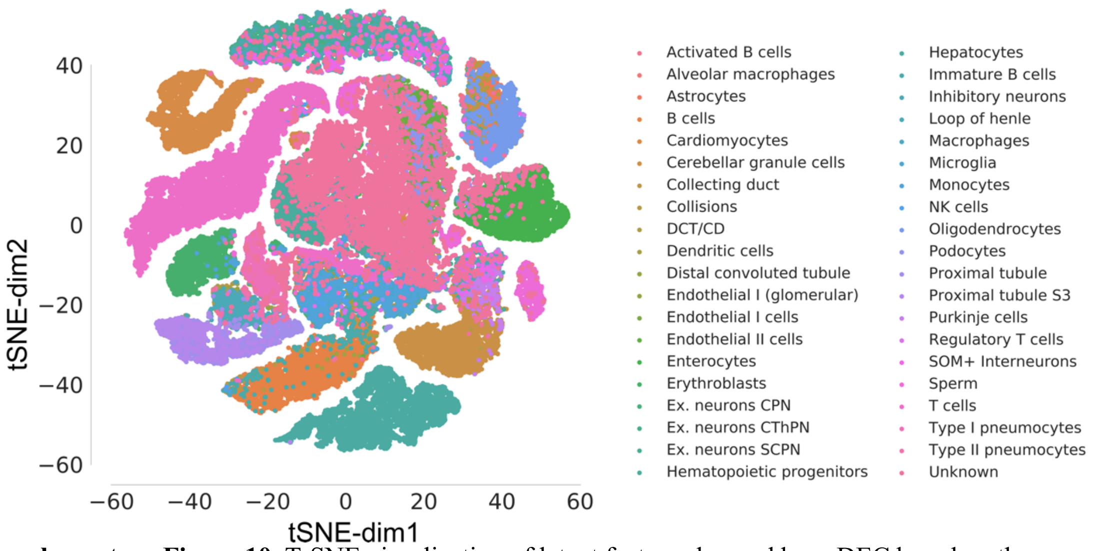


**Supplementary Figure 10.** T-SNE visualization of latent features learned by scDEC based on the Mouse atlas full dataset.

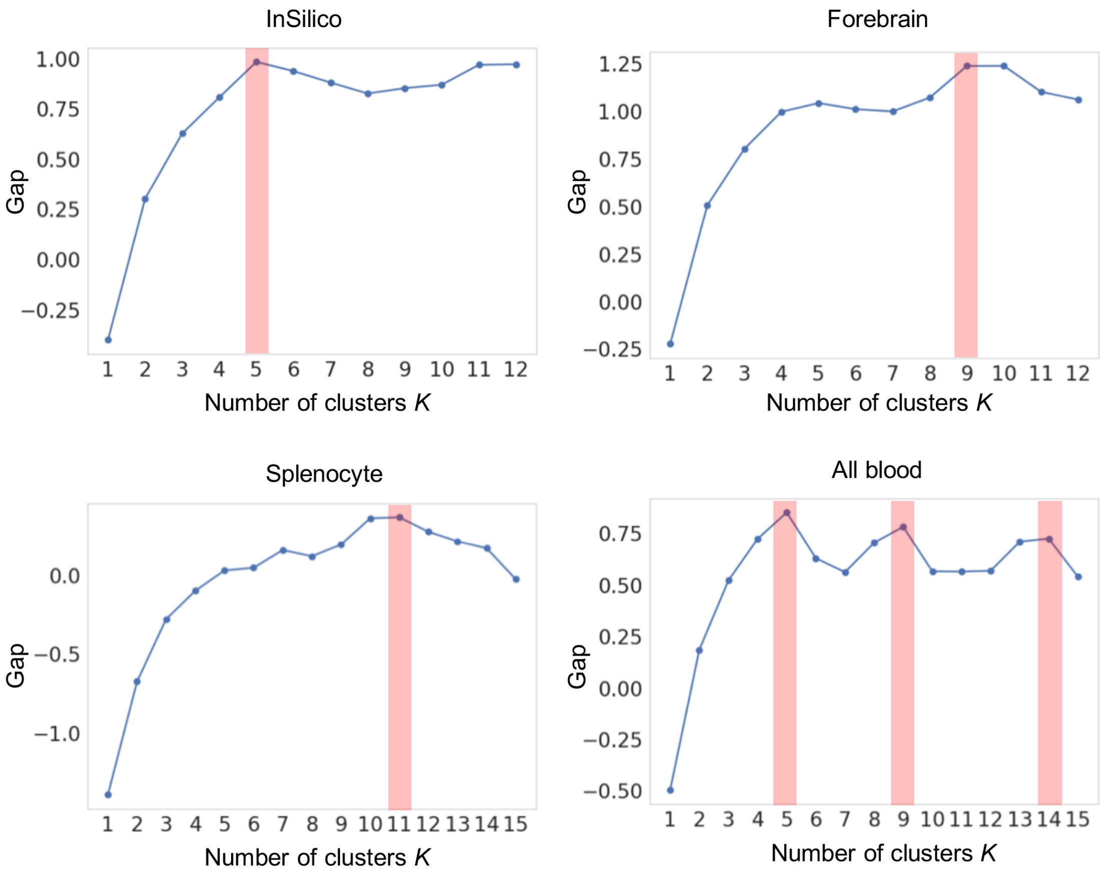


**Supplementary Figure 11.** The estimated number of clusters using gap statistic of four datasets used in this study.

### Supplementary Tables

**Supplementary Table 1**. The detailed hyperparameters of scDEC model. FC: fully-connected layer, ReLu: rectified linear unit, Identity: identity mapping (without non-linear activation function), Tanh: Tangent Hyperbolic function, Softmax: softmax function, batchnorm: batch normalization.

| **Generator** G | **Discriminator** $D_x$ |
|---|---|
| Inputs $\boldsymbol{z} \in \mathbb{R}^d$ and $\boldsymbol{c} \in \mathbb{R}^K$ | Input $\boldsymbol{x} \in \mathbb{R}^{20}$ or $\tilde{\boldsymbol{x}} \in \mathbb{R}^{20}$ |
| Concat($\boldsymbol{z}, \boldsymbol{c}$) | FC, 256, ReLu |
| (FC, 512, ReLu)$\times 10$ | FC, 256, batchnorm, Tanh |
| FC, 20, Identity | Fc, 1, Identity |
| Output $\tilde{\boldsymbol{x}} \in \mathbb{R}^{20}$ | Output $logit_x \in \mathbb{R}^1$ |
| **Generator** H | **Discriminator** $D_z$ |
| Input $\boldsymbol{x} \in \mathbb{R}^{20}$ | Input $\boldsymbol{z} \in \mathbb{R}^d$ or $\tilde{\boldsymbol{z}} \in \mathbb{R}^d$ |
| (FC, 256, ReLu)$\times 10$ | FC, 256, ReLu |
| FC, *d*, Identity and FC, *K*, Softmax | FC, 256, batchnorm, Tanh |
| Outputs $\tilde{\boldsymbol{z}} \in \mathbb{R}^d$ and $\tilde{\boldsymbol{c}} \in \mathbb{R}^K$ | Fc, 1, Identity |
| | Output $logit_z \in \mathbb{R}^1$ |

**Supplementary Table 2**. The summary of the five scATAC-seq datasets included in this study.

| Dataset | No. Of cells | No. of peaks | No. of cell types | Accession |
|---|---|---|---|---|
| InSilico | 1377 | 68069 | 6 | GEO:GSE65360 |
| ForeBrain | 2088 | 140102 | 8 | GEO:GSE100033 |
| Splenocyte | 3166 | 77453 | 12 | ArrayExpress:E-MTAB-6714 |
| All blood | 2034 | 455057 | 13 | GEO:GSE96772 |
| Mouse atlas | 81173 | 436206 | 30 | http://atlas.gs.washington.edu/mouse-atac |

**Supplementary Table 3**. The information of enriched motif discovered by scDEC. We used the Mann-Whitney U test for evaluating significance of motif enrichment with the alterative hypothesis that the chromVAR deviation score in one/multiple clusters are higher than the score in the rest of cells.

| Motif | Family | Enriched cluster | Enriched cell type | $p$-value |
|---|---|---|---|---|
| LBX2 | NK-related factors | 1 | Astrocyte | $1.04\times10^{-51}$ |
| EN1 | NK-related factors | 1 | Astrocyte | $6.14\times10^{-51}$ |
| BSX | NK-related factors | 1 | Astrocyte | $2.60\times10^{-50}$ |
| MSX2 | NK-related factors | 1 | Astrocyte | $2.33\times10^{-48}$ |
| MSX1 | NK-related factors | 1 | Astrocyte | $1.03\times10^{-46}$ |
| GBX1 | HOX-related factors | 1 | Astrocyte | $2.19\times10^{-45}$ |
| GBX2 | HOX-related factors | 1 | Astrocyte | $2.90\times10^{-45}$ |
| EN2 | NK-related factors | 1 | Astrocyte | $7.95\times10^{-45}$ |
| NOTO | NK-related factors | 1 | Astrocyte | $2.04\times10^{-44}$ |
| SHOX | Paired-related HD factors | 1 | Astrocyte | $3.59\times10^{-44}$ |
| NEUROD2 | Tal-related factors | 2-4 | Excitatory neuron | $4.50\times10^{-239}$ |
| OLIG1 | Tal-related factors | 2-4 | Excitatory neuron | $9.32\times10^{-239}$ |
| BHLHE22 | Tal-related factors | 2-4 | Excitatory neuron | $4.26\times10^{-236}$ |
| BHLHE23 | Tal-related factors | 2-4 | Excitatory neuron | $1.60\times10^{-233}$ |
| OLIG3 | Tal-related factors | 2-4 | Excitatory neuron | $1.08\times10^{-232}$ |
| OLIG2 | Tal-related factors | 2-4 | Excitatory neuron | $3.81\times10^{-231}$ |
| NEUROG2 | Tal-related factors | 2-4 | Excitatory neuron | $3.15\times10^{-225}$ |
| TAL1::TCF3 | Tal-related factors | 2-4 | Excitatory neuron | $2.49\times10^{-205}$ |
| MEIS3 | TALE-type homeodomain factors | 5 | Inhibitory neuron 1 | $1.44\times10^{-61}$ |
| MEIS1 | TALE-type homeodomain factors | 5 | Inhibitory neuron 1 | $6.68\times10^{-59}$ |
| ZEB1 | HD-ZF factors | 5 | Inhibitory neuron 1 | $1.13\times10^{-58}$ |
| MEIS2 | TALE-type homeodomain factors | 5 | Inhibitory neuron 1 | $1.32\times10^{-58}$ |
| DBP | C/EBP-related | 5 | Inhibitory neuron 1 | $5.80\times10^{-54}$ |
| TEF | TEF-1-related factors | 5 | Inhibitory neuron 1 | $7.01\times10^{-48}$ |
| HLF | C/EBP-related | 5 | Inhibitory neuron 1 | $1.35\times10^{-46}$ |
| TCF4 | E2A-related factors | 5 | Inhibitory neuron 1 | $2.36\times10^{-46}$ |
| NFIL3 | C/EBP-related | 5 | Inhibitory neuron 1 | $4.32\times10^{-46}$ |
| TCF3 | E2A-related factors | 5 | Inhibitory neuron 1 | $1.63\times10^{-44}$ |
| MEOX2 | HOX-related factors | 6 | Inhibitory neuron 2 | $9.36\times10^{-133}$ |
| VAX1 | NK-related factors | 6 | Inhibitory neuron 2 | $2.84\times10^{-126}$ |
| VAX2 | NK-related factors | 6 | Inhibitory neuron 2 | $2.84\times10^{-126}$ |

| | | | | |
|---|---|---|---|---|
| LMX1B | HD-LIM factors | 6 | Inhibitory neuron 2 | $9.97\times10^{-121}$ |
| MEOX1 | HOX-related factors | 6 | Inhibitory neuron 2 | $6.23\times10^{-116}$ |
| HOXB2 | HOX-related factors | 6 | Inhibitory neuron 2 | $1.16\times10^{-114}$ |
| HOXA2 | HOX-related factors | 6 | Inhibitory neuron 2 | $8.29\times10^{-112}$ |
| HOXB3 | HOX-related factors | 6 | Inhibitory neuron 2 | $2.52\times10^{-108}$ |
| NKX6-2 | NK-related factors | 6 | Inhibitory neuron 2 | $4.23\times10^{-107}$ |
| NKX6-1 | NK-related factors | 6 | Inhibitory neuron 2 | $2.23\times10^{-105}$ |
| SPIC | Ets-related factors | 7 | Microglia | $2.02\times10^{-73}$ |
| SPI1 | Ets-related factors | 7 | Microglia | $2.09\times10^{-73}$ |
| ETV6 | Ets-related factors | 7 | Microglia | $9.91\times10^{-72}$ |
| ELF5 | Ets-related factors | 7 | Microglia | $1.86\times10^{-71}$ |
| ELK1 | Ets-related factors | 7 | Microglia | $1.87\times10^{-71}$ |
| ETS1 | Ets-related factors | 7 | Microglia | $1.20\times10^{-66}$ |
| ERF | Ets-related factors | 7 | Microglia | $3.45\times10^{-66}$ |
| EHF | Ets-related factors | 7 | Microglia | $4.43\times10^{-66}$ |
| ELF4 | Ets-related factors | 7 | Microglia | $1.18\times10^{-65}$ |
| ELF3 | Ets-related factors | 7 | Microglia | $1.87\times10^{-65}$ |
| SOX9 | SOX-related factors | 8 | Oligodendrocyte | $3.81\times10^{-137}$ |
| JUN | Jun-related factors | 8 | Oligodendrocyte | $5.38\times10^{-105}$ |
| SRY | SOX-related factors | 8 | Oligodendrocyte | $1.77\times10^{-96}$ |
| JUND(var.2) | Jun-related factors | 8 | Oligodendrocyte | $7.08\times10^{-93}$ |
| HNF4G | RXR-related receptors | 8 | Oligodendrocyte | $2.01\times10^{-91}$ |

**Supplementary Table 4**. We have applied both Welch's t test and Mann-Whitney U test to verify the superiority of our method for data generation compared to interpolating PCA reduced data. As for the former, we test whether the two samples have the equal means. As for the later, we test with the alternative hypothesis that the PCCs of our method for the different cell have a positive shift when compared with those of a baseline.

| | Pearson r | Spearman r |
|---|---|---|
| Welch's t test | $9.97\times10^{-21}$ | $6.80\times10^{-7}$ |
| Mann-Whitney U test | $1.28\times10^{-16}$ | $4.40\times10^{-8}$ |

**Supplementary Table 5**. The training of scDEC model. The default settings in all the experiments are as follows. We use Adam optimizer for gradient descent and parameters updating. $\alpha = 0.002$ , $\mathrm{m} = 32$, the default parameters for Adam optimizer were used.

---

**Algorithm** Training procedure of scDEC

---

**Require**: $\theta_g^0$ and $\theta_h^0$ for initial parameters of G and H network, $\theta^0 = (\theta_g^0, \theta_h^0)$, $\omega_{dx}^0$ and $\omega_{dz}^0$ for initial parameters of $D_x$ and $D_z$ network, $\omega^0 = (\omega_{dx}^0, \omega_{dz}^0)$, batch size $m$ and learning rate $\alpha$.

**While** $\theta, \omega$ have not converged, **do**

  **For** $\mathrm{t} = 1, \dots, \mathrm{n}_d$, **do**

    **For** $\mathrm{i} = 1, \dots, \mathrm{m}$, **do**

      Sample data $\boldsymbol{x} \sim \mathcal{P}_{\boldsymbol{x}}$ , latent variable $\boldsymbol{z} \sim \mathcal{P}_{\boldsymbol{z}}, \boldsymbol{c} \sim \mathcal{P}_{\boldsymbol{c}}$ and a random number $\zeta, \eta \sim \mathrm{U}(0,1)$.

      $\tilde{\boldsymbol{x}} \leftarrow \mathrm{G}(\mathbf{z}, \mathbf{c})$

      $\tilde{\boldsymbol{z}}, \tilde{\boldsymbol{c}} \leftarrow \mathrm{H}(\boldsymbol{x})$

      $\hat{\boldsymbol{x}} \leftarrow \zeta \boldsymbol{x} + (1 - \zeta)\tilde{\boldsymbol{x}}$

      $\bar{\mathbf{z}} \leftarrow \eta \boldsymbol{z} + (1 - \eta)\tilde{\boldsymbol{z}}$

      $L_{dx}^{(i)} \leftarrow \mathrm{D}_x(\tilde{\boldsymbol{x}}) - \mathrm{D}_x(\boldsymbol{x}) + \lambda(||\nabla_{\hat{\boldsymbol{x}}} \mathrm{D}_x(\hat{\boldsymbol{x}})||_2 - 1)^2$

      $L_{dz}^{(i)} \leftarrow \mathrm{D}_z(\tilde{\boldsymbol{z}}) - \mathrm{D}_z(\boldsymbol{z}) + \lambda(||\nabla_{\bar{\boldsymbol{z}}} \mathrm{D}_z(\bar{\boldsymbol{z}})||_2 - 1)^2$

      $L_d^{(i)} = L_{dx}^{(i)} + L_{dz}^{(i)}$

    **End**

    $\omega \leftarrow \omega + \alpha \cdot \mathrm{Adam}(\nabla_\omega \frac{1}{m} \sum_{i=1}^{m} L_d^{(i)}, \omega)$

  **End**

  **For** $\mathrm{i} = 1, \dots, \mathrm{m}$, **do**

    Sample data $\boldsymbol{x} \sim \mathcal{P}_{\boldsymbol{x}}$ , latent variable $\boldsymbol{z} \sim \mathcal{P}_{\boldsymbol{z}}, \boldsymbol{c} \sim \mathcal{P}_{\boldsymbol{c}}$.

    $\tilde{\boldsymbol{x}} \leftarrow \mathrm{G}(\mathbf{z}, \mathbf{c})$

    $\tilde{\boldsymbol{z}}, \tilde{\boldsymbol{c}} \leftarrow \mathrm{H}(\boldsymbol{x})$

    $L_{g,h}^{(i)} \leftarrow -\mathrm{D}_x(\boldsymbol{x}) - \mathrm{D}_z(\tilde{\boldsymbol{z}}) + 10(||\boldsymbol{x} - \mathrm{G}(\mathrm{H}(\boldsymbol{x}))||_2^2 + ||\boldsymbol{z} - \mathrm{H}(\mathrm{G}(z))||_2^2 + CE(\boldsymbol{c}, \tilde{\boldsymbol{c}}))$

  $\theta \leftarrow \theta + \alpha \cdot \mathrm{Adam}\left(\nabla_\theta \frac{1}{m} \sum_{i=1}^{m} L_{g,h}^{(i)}, \theta\right)$

**End while**

---
